# LACHESIS: real-time inference of evolutionary trajectories of malignant transformation from whole-genome sequencing data

**DOI:** 10.64898/2026.07.24.740074

**Authors:** Maximilia Eggle, Anand Mayakonda, Christoph Bartenhagen, Thomas Höfer, Frank Westermann, Verena Körber

## Abstract

Computational tools for phylogenetic inference of early tumor evolution in real time are currently lacking. We present LACHESIS, a standardized R package and Shiny app that times early and most recent common ancestors of individual tumors from whole-genome sequencing data. LACHESIS automates mutational signature-aware molecular clock modeling for trajectory reconstruction and evolution-based risk stratification. We validate its utility for childhood and adult malignancies, providing a broadly applicable pan-cancer workflow.

## Main

Early stages of spontaneous tumor development are usually obscured from direct observation. Existing tools reconstruct subclonal evolution from the most recent common ancestor (MRCA) using bulk or single-cell sequencing^1-7^, but do not address earlier stages of malignant transformation. Aneuploidy is a frequent hallmark of early, (pre-)malignant states and leaves distinct footprints in somatic variant allele frequencies (VAFs). Building on computational frameworks that use these footprints to time chromosomal copy number gains^8-10^, we present LACHESIS: a scalable, user-friendly R package and Shiny Web application for early tumor trajectory reconstruction in chronological time. LACHESIS provides a unified computational pipeline that accepts flexible variant-calling inputs, integrates mutational signature data, and features evolution-based risk stratification. We demonstrate its pan-cancer utility for neuroblastoma, paraganglioma, and glioblastoma.

### LACHESIS, a computational framework for evolutionary reconstruction from WGS data

LACHESIS interprets somatic mutation accumulation as a molecular clock to infer tumor evolutionary histories from the number and allelic configuration of single nucleotide variants (SNVs)^9^. First, LACHESIS quantifies the emergence of a tumor’s MRCA from the abundance of clonal SNVs, identified based on VAF (Fig. 1a). Second, LACHESIS times clonal driver mutations and copy number gains by further analyzing VAF distributions in affected regions: to this end, LACHESIS quantifies the abundance of SNVs on multiple chromosomal copies and thus acquired before the copy number variant (CNV) (Fig. 1a). Comparing the abundance of pre-CNV SNVs to the overall abundance of clonal SNVs, LACHESIS distinguishes whether clonal copy number gains occurred concomitant with a tumor’s MRCA or before, in an early common ancestor (ECA). Lastly, evolutionary timelines are translated into chronological time using either established mutation rates^9-13^ or a new modeling approach implemented in LACHESIS, which infers the pre-MRCA SNV accumulation rate by linear regression of MRCA mutation burden against age at diagnosis (Fig. 1b and Methods; Benedetto et al., submitted).

LACHESIS is compatible with SNV, CNV, ploidy and purity data generated by a wide range of bioinformatic workflows. It seamlessly integrates mutational signature information via a direct interface to MutationalPatterns^8^or by importing output from SigProfilerAssignment^14^. Finally, LACHESIS supports evolution-based risk stratification (Fig. 1c).

**Figure 1.**
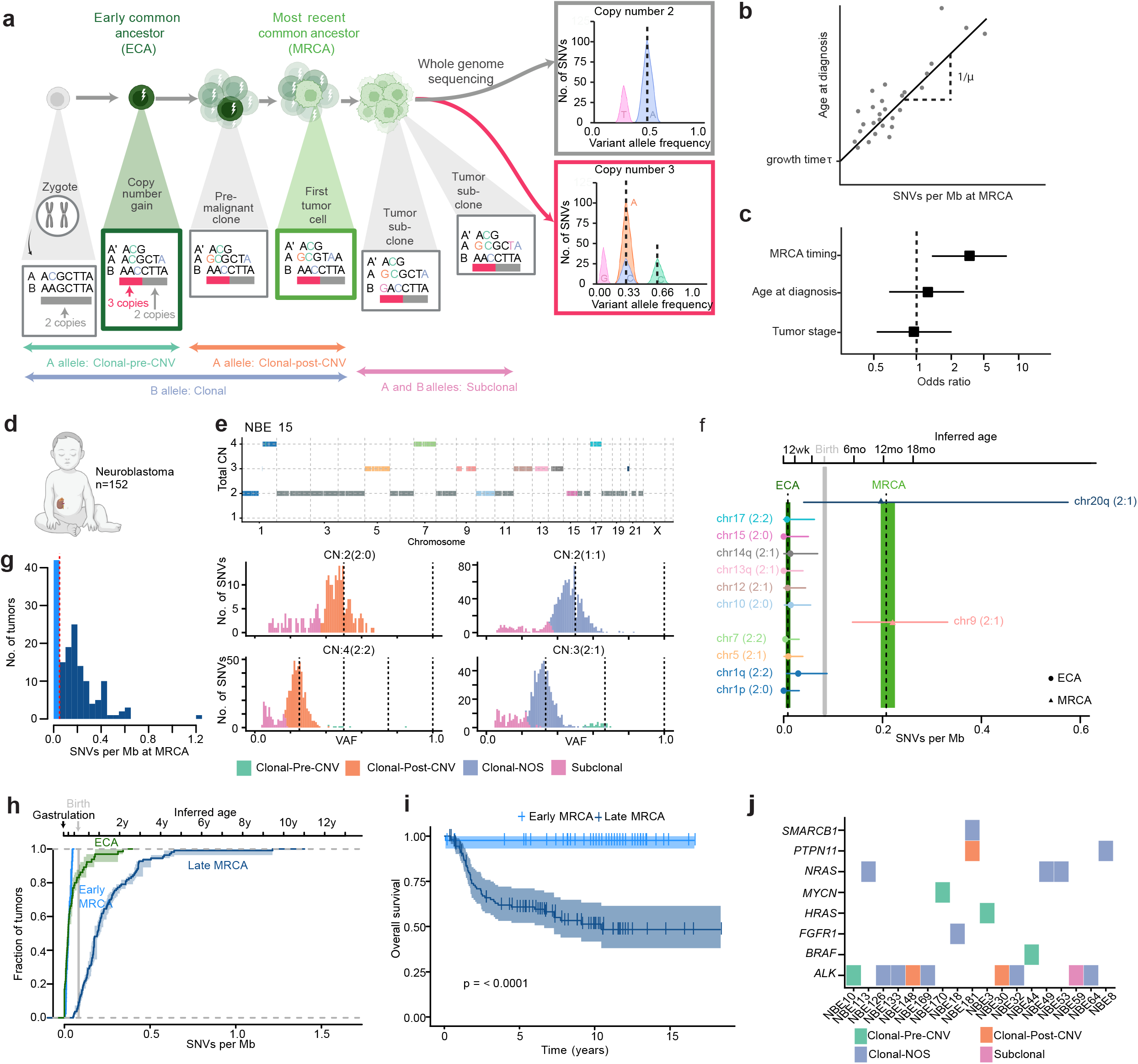
Reconstructing tumor evolution with LACHESIS. **a**, LACHESIS times tumor initiation using somatic mutations as molecular barcodes. Clonal somatic mutations are characteristic of the tumor’s most recent common ancestor (MRCA). In a diploid genome, they typically have heterozygous variant allele frequencies (VAF) of 0.5. Some tumors also acquire copy number variants (CNVs) during malignant transformation. CNVs can be acquired in the tumor’s MRCA, or before, in an early common ancestor (ECA). CNVs change the VAFs of somatic mutations present in the affected region and can therefore be identified as distinct clusters in the measured VAF histogram (right plots): mutations acquired before a CNV and on gained copies have higher VAF than mutations acquired after the gain or on non-gained alleles (compare green to orange and blue mutations). Finally, mutations acquired during tumor growth will be subclonal (pink). **b**, Scheme illustrating mutation rate estimation based on linear regression between age at diagnosis and the SNV burden in the tumor’s MRCA. **c**, Scheme illustrating the potential of LACHESIS for evolution-based risk stratification. **d**, LACHESIS is exemplified on a cohort of 152 neuroblastomas. **e**, Example output of LACHESIS, showing copy number profile and timing of mutations. Clonal Pre-CNV, mutation acquired before a copy number gain; clonal Post-CNV, mutation acquired after a copy number gain; Clonal-NOS, clonal not otherwise specified. **f**, Inferred evolutionary timeline for the example shown in c (dashed lines and shaded areas, mean and 95% confidence intervals (CIs) of estimated mutation densities at ECA and MRCA; grey area, estimated mutation density at birth; points and error bars, mean and 95% CIs for mutation densities at chromosomal gains). Inferred age is based on the cohort-derived mutation rate of 2.17 SNVs per day (95% CI: 1.52-3.77 SNVs per day; Extended Data Fig. 1a, b). **g**, Inferred mutation densities at MRCA for 152 neuroblastomas. **h**, Inferred mutation densities at ECA and MRCA for 152 neuroblastomas. Shown are mean and 95% CIs of the cumulative distribution. Inferred age as in f. **i**, Overall survival of 152 neuroblastomas stratified by MRCA time. Shown are mean and 95% confidence bounds according to a log-rank test (p = 0.00000237). Tumors with mutation densities at MRCA below 0.05 SNVs per Mb were classified as early-MRCA. **j**, Driver mutations, identified in the cohort of 152 neuroblastomas; Clonal Pre-CNV, mutation acquired before a copy number gain; clonal Post-CNV, mutation acquired after a copy number gain; Clonal-NOS, clonal not otherwise specified.

We validated LACHESIS using WGS data of 152 primary neuroblastoma samples the early evolution of which we had previously studied in detail^9^(Fig. 1d-j, Extended Data Fig. 1). A representative example of a high-risk neuroblastoma with several copy number gains and alternative lengthening of telomeres is shown in Fig. 1e and f. Here, LACHESIS estimates a mutational burden of approximately 0.2 SNVs/Mb in the tumor’s MRCA, corresponding to approximately 12 months of life, using the cohort-derived mutation rate of 2.17 SNVs per day (Extended Data Fig. 1a). Clonal copy number gains were acquired in a significantly earlier ECA, timed within the first 12 weeks of gestation.

Across the cohort, SNV densities at ECA and MRCA were comparable to prior analysis and estimated per-day mutation rates were similarly consistent^9^(Extended Data Fig. 1a-c). Moreover, inference of tumor evolution with LACHESIS was robust to diverse SNV and CNV callers (Extended Data Fig. 1d-f). In agreement with our prior results, mutation densities in the MRCA were bimodal across the cohort (Fig. 1g, h), and high mutation burden was strongly associated with poor survival, acquired telomere maintenance and an ECA in the first gestational trimester (Fig. 1h, i, Extended Data Fig. 2a). In a subset of high-risk cases, oncogenic mutations *ALK* p.R1275Q, *HRAS* p.Q61K, *BRAF* p.G469A and *MYCN* p.P44L occurred prior to the earliest chromosomal gains and were likely tumor initiating (Fig. 1j, Extended Data Fig. 2b-e). Thus, LACHESIS suggests that high-risk neuroblastomas date back to early fetal development and evolve via stepwise acquisition of copy number gains, telomere maintenance mechanisms and, in some cases, additional driver mutations.

### Evolutionary origin of paraganglioma

We next studied paragangliomas, neuroendocrine tumors with frequent germline predisposition, most commonly due to mutations in *SDHB*^15^. *SDHB* mutation is associated with increased risk of metastasis, making this subtype particularly relevant for improved clinical risk stratification^16^. We analyzed 43 primary *SDHB*-deficient paragangliomas from a recently published cohort^17^ (Fig. 2a). All tumors harbored copy number alterations involving Chromosome 1; eight cases had additional somatic driver mutations in *EPAS1, ATRX, MYCN* or the *TERT* promoter (Fig. 2b).

**Figure 2.**
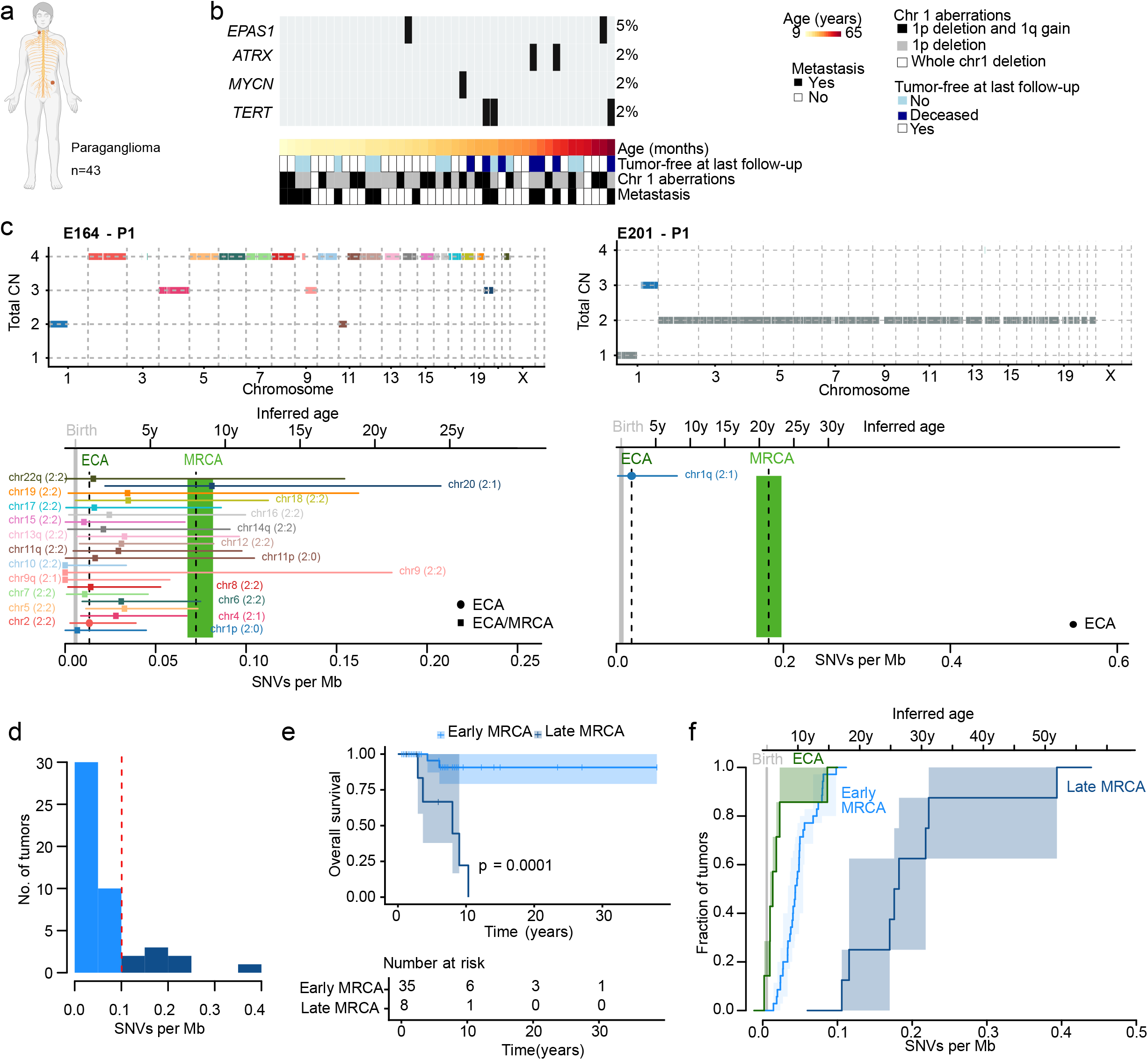
Diverging evolutionary trajectories in *SDHB*-mutant paragangliomas. **a**, Cohort overview. **b**, Mutational profile and associated clinical characteristics. **c**, Copy-number profiles and inferred evolutionary timelines of two representative tumors. Dashed lines and shaded areas, mean and 95% CIs of estimated mutation densities at ECA and MRCA; grey area, estimated mutation density at birth; points and error bars, mean and 95% CI of mutation densities at chromosomal gains. Inferred age based on cohort-derived mutation rate of 0.15 SNVs per day (95% CI: 0.10-0.28 SNVs/day; Extended Data Fig. 3c). **d**, Inferred mutation densities at MRCA for 43 paragangliomas. A threshold of 0.1 SNVs/Mb (red line) separates early- and late-MRCA tumors with distinct survival. **e**, Overall survival of 43 paragangliomas stratified by MRCA time (at a threshold of 0.1 SNVs/Mb). Shown are mean and 95% confidence bounds according to a log-rank test (P= 0.0001304741). **f**, Inferred mutation densities at ECA and MRCA for 43 paragangliomas. Shown are mean and 95% CIs intervals of the cumulative distribution. Inferred age as in e.

Applying LACHESIS to SNVs assigned to the clock-like mutational signatures SBS1, SBS5 and SBS40, we identified footprints of evolution before actual tumor growth. Among 24 cases with clonal copy number gains suitable for analysis (copy number ≤ 4, minimal segment size, 10^7^bp), seven showed evidence of an ECA, characterized by Chromosome 1q gain (n=4), triploidization (n=1) or tetraploidization (n=2) (Fig. 2c and Supplementary Tables 1 and 2). The allelic configuration in two tetraploid tumors allowed further resolution of the sequence of chromosomal events. Specifically, an A:B allelic configuration of 2:0 indicated that loss of Chromosome 1p preceded whole-genome duplication and hence the onset of tumor growth (Fig. 2c).

Mutation densities in the tumors’ MRCA varied widely (0.01-0.39 SNVs/Mb; Fig. 2d), suggesting substantial heterogeneity in the timing of tumor initiation. Consistent with that observation, MRCA mutation density correlated with age at diagnosis (Extended Data Fig. 3a). We therefore asked whether MRCA timing was associated with clinical outcome, as previously reported for neuroblastoma^9^. Indeed, paragangliomas with a late MRCA (≥0.1 SNVs/Mb) had significantly worse overall and metastasis-free survival than those with an early MRCA (<0.1 SNVs/Mb; p = 0.0001, Fig. 2e; Extended Data Fig. 3b).

Finally, we assessed the timing of ECA and MRCA across risk groups. LACHESIS estimated that the paraganglioma cell of origin accumulates on average 0.15 SNVs per day, substantially less than observed for neuroblastoma (Extended Data Fig. 1a; Extended Data Fig. 3c). This mutation rate suggests that both ECA and early MRCAs occured during childhood and adolescence, whereas the majority of late MRCAs occurred in early adulthood (Fig. 2f). These findings suggest that chromosomal gains in paraganglioma arise within a confined developmental time window, while the onset of malignant expansion spans a much broader period and defines clinical outcome.

Collectively, evolutionary analysis with LACHESIS suggests that the trajectories of low-risk and high-risk paragangliomas diverge early, potentially opening up new avenues for risk stratification.

### Evolutionary origin of IDH^WT^glioblastoma

To test whether LACHESIS reconstructs evolutionary timelines beyond adolescence, we assessed a cohort of 17 primary adult IDH^WT^ glioblastoma^18^ (Fig. 3a; Supplementary Table 3), again selecting for clock-like SNVs. A hallmark of IDH^WT^ glioblastoma is gain of Chromosome 7/7q. Using LACHESIS, we found that this gain occurs early, either as a solitary event (n=6) or in combination with additional chromosomal alterations, such as Chromosome 19 (n=6; Fig. 3b and c, Supplementary Tables 4 and 5).

**Figure 3.**
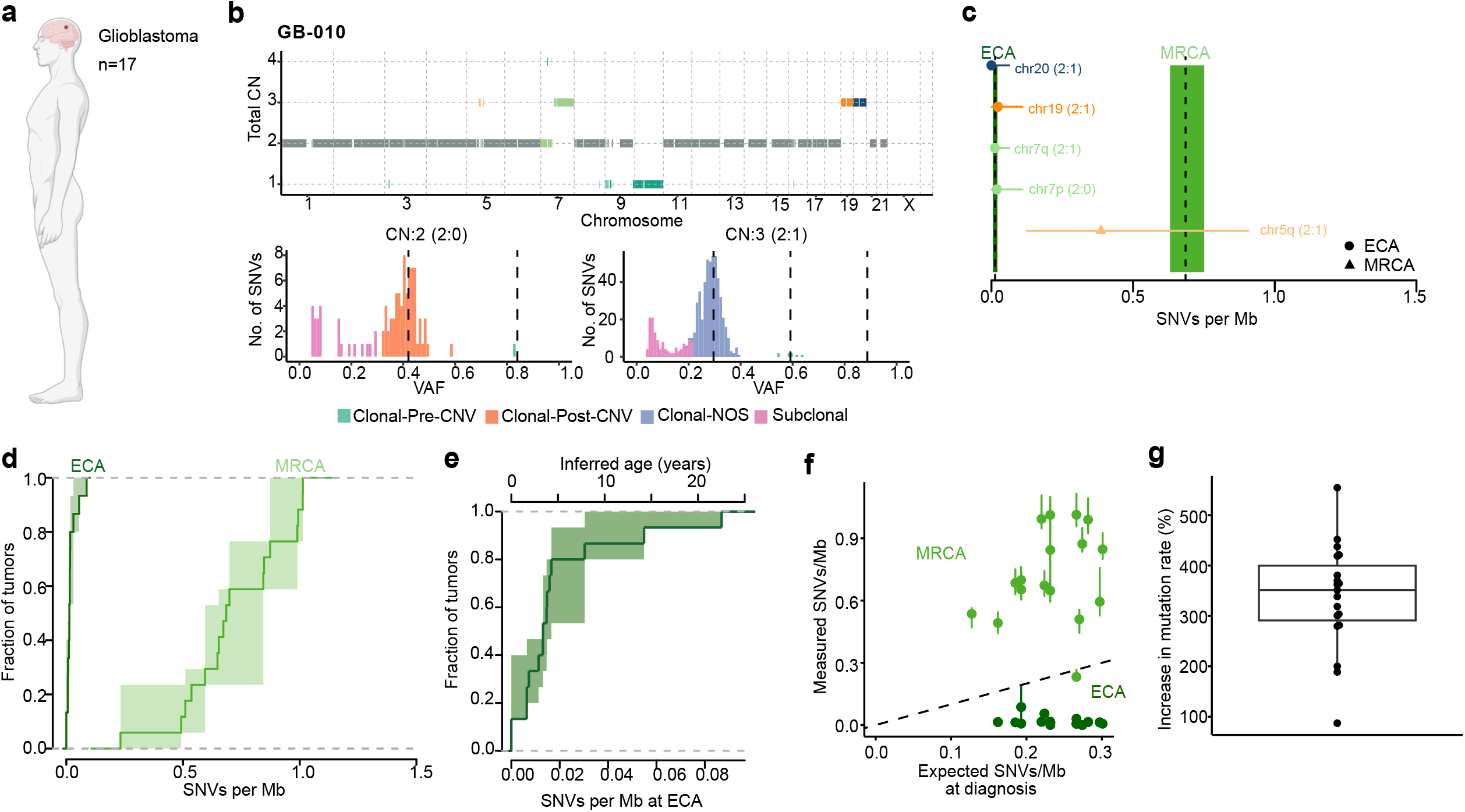
Early onset of glioblastoma evolution. **a**, Cohort overview. **b**, LACHESIS output for one representative example. Clonal Pre-CNV, clonal mutation acquired before a copy number gain; Clonal Post-CNV, clonal mutation acquired after a copy number gain; Clonal-NOS, clonal mutation, not otherwise specified. **c**, Inferred evolutionary timeline of the tumor shown in b. Dashed lines and shaded areas, mean and 95% CIs of estimated mutation densities at ECA and MRCA. Point and error bars, mean and 95% CIs for mutation densities at chromosomal gains. **d**, Mutation densities at ECA and MRCA across the cohort (n=17). Shown are mean and 95% CIs intervals of the cumulative distribution. **e**, Mutation densities at ECA (n=14) and inferred chronological time using physiological mutation rates in human oligodendrocytes^12^. Shown are mean and 95% CIs intervals of the cumulative distribution. **f**, Expected mutation densities at diagnosis, calculated from patient age at diagnosis and physiological mutation rates in human oligodendrocytes ^12^, compared with observed mutation densities (n=17). **g**, Ratio between observed and expected mutation densities described in f, reflecting a potential increase in mutation rate during glioblastoma evolution.

Mutation densities at ECA were uniformly very low (range 0-0.057 SNVs/Mb, Fig. 3d), suggesting that aneuploidies in glioblastoma are acquired early in life, despite the disease usually being diagnosed in midlife or later^19^. The confined number of cases in this cohort did not allow us to estimate SNV rates in glial cells *de novo*. We hence used published SNV accumulation rates in healthy human oligodendrocytes^12^ (0.07 SNVs/day for SBS1 and 5), and inferred that ECAs arose in childhood or adolescence (Fig. 3e). This early evolutionary origin raises the question whether glioblastomas undergo a decade-long pre-malignant phase before the onset of rapid tumor growth, or instead grow more slowly than previously assumed. To address this question, we translated mutation densities at MRCA into real time, again using mutation rates from healthy oligodendrocytes. This analysis yielded implausibly late MRCAs, with a mean age of 187 years (range 59-266 years) and thus incompatible with the age at diagnosis (Fig. 3f). Specifically, the observed ages at diagnosis require at least a 3.5-fold increase in mutation rate relative to healthy oligodendrocytes, strongly indicating that mutation accumulation accelerated during glioblastoma evolution (Fig. 3g). Together, these findings suggest that clonal trajectories of IDH-wildtype glioblastoma originate in early childhood, with mutation accumulation accelerating during malignant transformation.

## Discussion

Here we present LACHESIS, a robust, broadly applicable R/Bioconductor package and Shiny application to reconstruct early cancer evolution in chronological time. In contrast to alternative approaches that time copy number gains relative to a tumor’s MRCA^8, 20^, LACHESIS leverages absolute SNV counts and translates them into chronological time, while explicitly accounting for the statistical uncertainty inherent to small mutational burdens. Moreover, LACHESIS seamlessly incorporates multi-caller inputs, mutational signature information and clinical data.

Applying LACHESIS to three cancers, neuroblastoma, paraganglioma and glioblastoma, we demonstrate its applicability across pediatric, adult, sporadic and hereditary tumor entities. Notably, we find that the evolutionary trajectories of low-risk and high-risk neuroblastomas and paragangliomas diverge early, offering new avenues for improved risk stratification. Moreover, we find a striking latency between malignant transformation and clinical diagnosis: pediatric neuroblastomas date back to early gestation, and adult glioblastomas to childhood, opening up potential opportunities for early screening and intervention.

Mutation rates in neuroblastoma were an order of magnitude higher than those in paraganglioma, possibly reflecting differences between their putative cells of origin, sympathoblasts and chromaffin cells. In glioblastoma, we, moreover, find that mutation rates accelerate over the course of malignant transformation. Metabolic changes, replication stress or inflammatory signaling within the tumor microenvironment could contribute to this observation. Alternatively, early mutations could allow rapidly dividing progenitor cells to become self-renewing, leading to a replication-associated increase in mutation rate. Such scenarios have been described in acute myeloid leukemia^19^ and it will be fascinating to see whether they apply to solid tumors, too.

The observed acceleration in mutation rate highlights a core limitation of LACHESIS: real-time estimates rely on the assumption that mutation rates remain (approximately) constant over time and should therefore be interpreted with caution. However, we expect that advancements in measuring mutation rates across normal and diseased tissues^12, 13, 21, 22^ will allow incorporation of more detailed information into LACHESIS in the future. Additional technical constraints include a requirement of large copy number gains (>100 Mb) with tractable states (≤4), and the fact that the absence of an ECA does not preclude unresolved early evolutionary states.

Overall, LACHESIS democratizes evolutionary timing, providing an automated, scalable framework to decipher the earliest stages of human carcinogenesis, to guide the engineering of temporally accurate tumor models, and to advance early intervention and clinical risk stratification.

## Methods

### The model

The statistical model underlying LACHESIS consists of four major steps. First, a binomial mixture model is employed to quantify the number of clonal and subclonal SNVs. Second, the mutation burden at the onset of tumor growth from its MRCA is estimated. Third, negative binomial models are used to time the acquisition of copy number gains. Finally, the mutation burden can be converted into chronological time by relating age at diagnosis with mutation burden at MRCA across cohorts. The statistical model will be briefly laid out in the following and has been described in detail previously^9^(Benedetto et al., submitted).

#### 1. Quantifying the number of clonal SNVs

LACHESIS considers copy number states with a total copy number *TCN = A + B*. Depending on the allelic configuration and the timing of mutation acquisition, SNVs may be present on a single copy, on all copies of the A allele or on all copies of the B allele. Let *S={1, A, B}* denote the set of distinct allelic states, with duplicate values removed when *A=1, B=1* or *A=B*. We define the mixture proportion vector 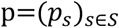 constrained to the simplex *Δ*^|*S*|-1^, as the relative proportion of SNVs assigned to each clonal allelic state. The maximum likelihood estimate for p is

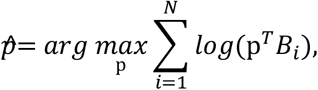

where *N* is the number of SNVs and 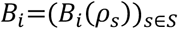 is the vector of binomial likelihoods for SNV *i* under each allelic state. Specifically, 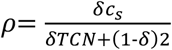 where *δ* is the purity of the sample and *c*_*S*_ *∈ {1, A, B}* is the copy number associated with state *s*.

#### 2. Quantifying mutation burden at MRCA

The timing of a tumor’s MRCA is quantified as a copy-number-normalized clonal mutation count. To account for differences in copy-number state, LACHESIS estimates, for each segment, the mutation burden expected in a single haploid lineage as:

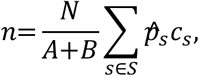

Where *N* denotes the number of clonal SNVs assigned to the segment. Subsequently, the mutation burden in the tumor’s MRCA is calculated as the sum of the copy-number-normalized mutation burdens across all genomic segments

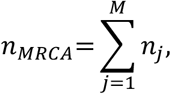

where *M* denotes the number of analyzed segments.

LACHESIS tests for each copy number gain of adequate length (≥10^7^bp) and copy number (A>1 or B>1, and TCN≤4) whether it occurred concomitant with the tumor’s MRCA. Under the null hypothesis that a gain occurred concomitantly with the MRCA, the number of mutations carried on the gained allele is expected to be proportional to the segment length and the genome-wide MRCA burden. To account for overdispersion of the mutation rate along the genome, LACHESIS models the null hypothesis with a negative binomial distribution, such that

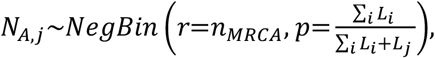

where *N*_*A,j*_ is the number of mutations on *A* copies on segment *j*, and *L*_*j*_ is the length of that segment (copy number states of the B allele are modeled in analogy). Under this parametrization, the expected mutation count is 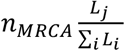 Segments with a Holm-corrected p value below 0.01 are considered inconsistent with a gain arising concomitantly with the MRCA. Such segments were classified as pre-MRCA if 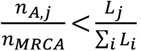 and post-MRCA otherwise.

#### 3. Quantifying mutation burden at ECA

In a subsequent step, LACHESIS aggregates all segments classified as pre-MRCA to estimate the mutation burden of the ECA:

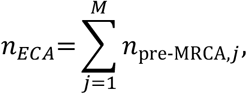

where 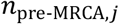 is the mutation burden associated with a gained segment classified as pre-MRCA. This aggregated burden is then used as the null expectation in a segment-wise negative binomial test to assess whether each segment is consistent with having arisen concomitantly with the ECA. As before, segments with a Holm-adjusted p value below 0.01 are considered incompatible with an origin in a common ECA.

#### 4. Translating SNV burden into chronological time

Following Benedetto et al. (submitted), LACHESIS models age at diagnosis as the time between gastrulation (the earliest time point from which somatic mutations can be counted using WGS) and the tumor’s MRCA, 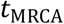 plus a tumor’s growth time, τ. Assuming a constant mutation rate, *μ*, before the tumor’s MRCA, the expected number of mutations in the MRCA is

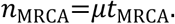

Accordingly, age at diagnosis can be expressed as

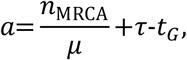

where we express age at diagnosis as the sum between patient age post birth, and the duration of gestation post gastrulation, *t*_*G*_. Assuming that τ is, on average, comparable between individuals, we can estimate *μ* using linear regression between the age at diagnosis and the SNV burden at MRCA across a cohort of tumors.

### Software functionality

LACHESIS is implemented as an R package and published as a continuously maintained package on Bioconductor. LACHESIS can be run by utilizing its sub-functions individually, or alternatively, by performing a streamlined workflow using the wrapper function `LACHESIS`.

The wrapper function requires as input estimates for tumor ploidy and purity, and the names and locations of the vcf files containing somatic SNV calls and bed files containing somatic copy number information. LACHESIS is designed to flexibly accept input from a plethora of variant callers; an overview of variant callers we have evaluated can be found in Supplementary Table 6. The following section outlines modalities associated with the different subfunctions. Full documentation is also provided in an accompanying vignette.

#### Importing data

The functions `readCNV` and `readVCF` are evoked by the LACHESIS main function to read in SNV calls and somatic copy number information. The two functions perform several quality checks on the input files, and provide reformatted, standardized output. `readCNV` further simplifies the copy number information by merging adjacent segments with equal copy number if they are separated by less than 10^5^ bp. If possible, allele-specific copy number information should be provided. If this information is unavailable, LACHESIS assumes that a given copy number reflects the allelic configuration requiring the minimal number of genomic changes. For example, a total copy number of 2 would be assigned to 1 A allele and 1 B allele, whereas a total copy number of 3 would be assigned to 2 A alleles and 1 B allele. The function `nbImport` integrates CNV and SNV information into a single data table, where each variant is assigned to its corresponding copy number segment and status.

#### Inference of clonal evolution

The function `ClonalMutationCounter` estimates the number of clonal mutations associated with a single or multiple copies per genomic segment, where segments of equal copy number and allelic configuration are chromosome-wise merged, at a tolerance for gaps between segments that can be specified by the user. This step involves a Binomial mixture model to infer the relative sizes of clonal peaks in the measured VAF distribution from the number of reference and variant reads per mutant locus (see above). To render clonal mutation counts comparable across genomic segments with different allelic configuration, the counts are normalized by the function `normalizeCounts`, which returns the mutation burden expected in a single haploid lineage. LACHESIS provides two modalities to incorporate mutational signatures, controlled by the parameter “sig.assign”. A built-in interface to the R package MutationalPatterns^22^ assigns individual mutations to mutational signatures. Alternatively, users can choose to learn mutational signatures independently of LACHESIS by using SigProfiler, and providing the output to LACHESIS using the “sig.file” parameter. If mutational signatures are incorporated into the workflow, mutation timing can be restricted to specific signatures of interest.

Finally, function MRCA estimates mutation densities at chromosomal gains, at MRCA and, if present, ECA from the normalized clonal mutation counts. Optionally, mean and standard deviation of a false discovery rate for clonal mutations (e.g., due to incomplete tissue sampling) can be provided. In a first step, the mutation burden in the MRCA is estimated from the total, normalized clonal mutation count. 95% confidence intervals are estimated by bootstrap resampling of genomic segment (1,000 replicates). The function MRCA then tests whether individual chromosomal gains occurred concomitantly with the MRCA. To this end, mutation densities for each chromosomal gain are estimated (together with 95% confidence intervals according to a Chi2 distribution). To test whether a chromosomal gain occurred concomitantly with the MRCA, the distribution of mutations along the genome is modeled using a negative binomial distribution, thus accounting for overdispersion. Chromosomal gains with a mutational burden significantly lower than expected at the MRCA (Holm-adjusted p value below 0.01) are classified as candidates for a pre-malignant state that emerged from an ECA. To test this hypothesis, the mutation burden in the ECA is estimated from all those chromosomal gains, before another round of negative binomial tests is performed that assesses for each segment whether it is consistent with having arisen in the ECA. The final assignment of CNVs to ECA, MRCA or alternative states is provided together with the respective *p* values. Lastly, the function `estimateClonality` reports for each SNV whether it is subclonal or clonal; if the SNV falls on a gained region, it further classifies the timing of the SNV relative to the copy number gain.

#### Translating mutation time into chronological time

LACHESIS translates mutation time into chronological time by using either established mutation rates for a tissue of interest or by inferring mutation rates *de novo*. For *de novo* inference, LACHESIS fits a linear regression of age at diagnosis against SNV burden in the tumor’s MRCA across a cohort. This approach assumes that the age at diagnosis is determined by the time of MRCA emergence and the subsequent tumor growth period. If growth durations are comparable across tumors, variation in age at diagnosis primarily reflects variation in MRCA timing. Because MRCA time is proportional to the SNV burden at MRCA, with the inverse mutation rate as the proportionality constant, the regression slope provides an estimate of the mutation rate.

#### Survival analysis based on tumor evolution

For entities, where MRCA timing holds predictive value, the downstream function `classifyLACHESIS` facilitates the use of LACHESIS in translational settings. Specifically, this function categorizes tumors into groups with early or late MRCA, using either a predefined thresholds of by searching for associations between MRCA time and clinical outcome. The output can be visualized using the function `plotSurvival`.

#### Visualization functions

Plotting functions are provided to visualize the output of LACHESIS. The function `plotNB` generates copy number profiles and VAF histograms, stratified by clonality or mutational signature, while the function `plotMutationDensities` visualizes inferred evolutionary trajectories. To compare evolutionary timings within cohorts, LACHESIS provides the functions `plotDiseaseTrajectories` and `plotLACHESIS`, which visualize the distribution of mutation densities at ECA and MRCA across tumors, and the function `plotClinicalCorrelations`, which compares SNV densities at ECA and MRCA with clinical parameters.

### Human datasets

We analyzed previously published WGS data from 152 neuroblastomas^9, 23, 24^, 43 paragangliomas^17^ and 17 glioblastomas^18^. The 45 neuroblastoma samples from Peifer et al. were analysed with different SNV calling pipelines (SNVCallingWorkflow, Strelka-modified version of Peiflyne) and used for benchmarking. Somatic variant calls and purity/ploidy estimates were obtained from the original studies, which used AceSeq, SNVCallingWorkflow, Strelka-modified version of Peiflyne for neuroblastoma and glioblastoma, and PURPLE and bcbio for paranganglioma. Samples were included if they met the following criteria:

1. Estimated tumor purity >30%;
2. Tumor mutational burden <10 SNVs/Mb, thus focusing on cases without hypermutation due to defective DNA repair;
3. Reported tumor purity and ploidy estimates verified by concordance between expected and observed VAFs.

### Mutational signature analysis

Mutational signatures were assigned using SigProfilerAssignment (version 1.1.3) and COSMIC v3.5 reference signatures. Analyses using clock-like mutational signatures were restricted to SBS1, SBS5 and SBS40.

### LACHESIS analysis

Analyses were performed with LACHESIS v1.1.1, available on https://bioconductor.org/packages/devel/bioc/html/LACHESIS.html. In all cohorts, the minimum depth for a variant to be considered was set to 0, in order to retain all genomic regions. For the neuroblastoma cohort, we used a clonal mutation false-positive rate of 0.15 and a standard deviation of 0.05, reflecting incomplete sampling of the resected tumor by WGS, as previously described^9^ All remaining parameters were left at their default values as described in Methods-Main software functions.

### Driver mutation analysis

Driver mutations were compiled from cancer hotspot database (https://www.cancerhotspots.org/#/home)^25^ and are available on our GitHub repository (LACHESIS/inst/extdata/cancer_hotspots_v2_GRCh37_adapted.tsv).

### Use of generative AI

The authors used ChatGPT (OpenAI, GPT-5.5) to assist with language editing and code optimization during software development. All AI-generated suggestions were critically reviewed, tested and modified as appropriate by the authors. The authors developed and verified all scientific content, analyses, interpretations and conclusions, and take full responsibility for the final manuscript and software.

### Statistics and reproducibility

LACHESIS was developed on R v.4.3.1, R v4.4.1 and R v4.5.1. All statistical tests are described in the main text, methods section and in the Figure legends.

## Data availability

WGS data were part of previously published studies.^9, 23, 24, 17^,^18^. WGS data from these studies are deposited at the European Genome-Phenome Archive (https://www.ebi.ac.uk/ega/) under accession nos. EGAS00001004349, EGAS00001001308, EGAS00001004990, EGAS0000100653, EGAS50000000346 and EGAS00001003184.

## Code availability

LACHESIS is available on GitHub (https://github.com/VerenaK90/LACHESIS), Bioconductor (DOI: 10.18129/B9.bioc.LACHESIS; most recent version: https://bioconductor.org/packages/devel/bioc/html/LACHESIS.html) and as a ShinyApp on https://lachesis.dkfz.de.

## Figure legends

**Extended Data Figure 1.**
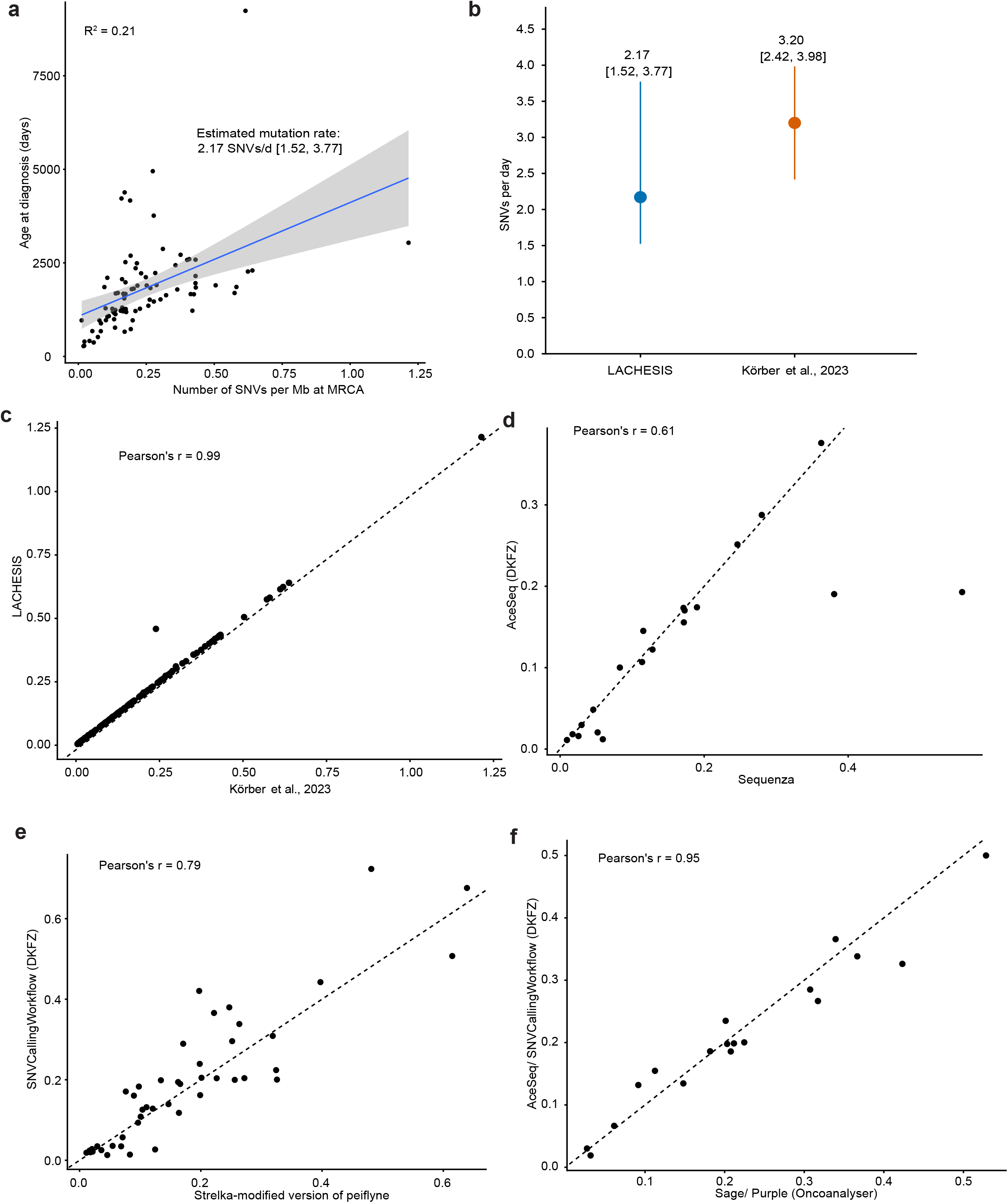
Performance of LACHESIS across bioinformatic pipelines. **a**, Linear regression relating age at diagnosis to SNV burden in the tumor’s MRCA across 87 high-risk neuroblastomas (points, individual tumors; solid line, fitted linear regression line (R^2^ = 0.21); grey shaded area, 95% confidence interval of fitted mean). **b**, Comparison of the SNV accumulation rate estimated by LACHESIS with the estimate reported previously^9^(points, estimated mutation rate in SNVs per day, error bars, 95% confidence intervals). No significant difference was observed between the two estimates (two-sided Z-test, z=-1.47, P=0.141). **c**, Mutation densities at MRCA for 152 neuroblastomas estimated previously^9^ and with LACHESIS. **d**, Mutation densities at MRCA for 20 neuroblastomas where copy-number calls from both AceSeq and Sequenza were available. **e**, Mutation densities at MRCA for 45 neuroblastomas^24^ where SNV calls from both SNVCallingWorkflow and Strelka-modified version of Peiflyne^26^ were available. **f**, Mutation densities at MRCA for 18 neuroblastomas where CNV and SNV calls from both Oncoanalyser pipeline (Sage/Purple) and the DKFZ pipeline (AceSeq/SNVCallingWorkflow) were available.

**Extended Data Figure 2.**
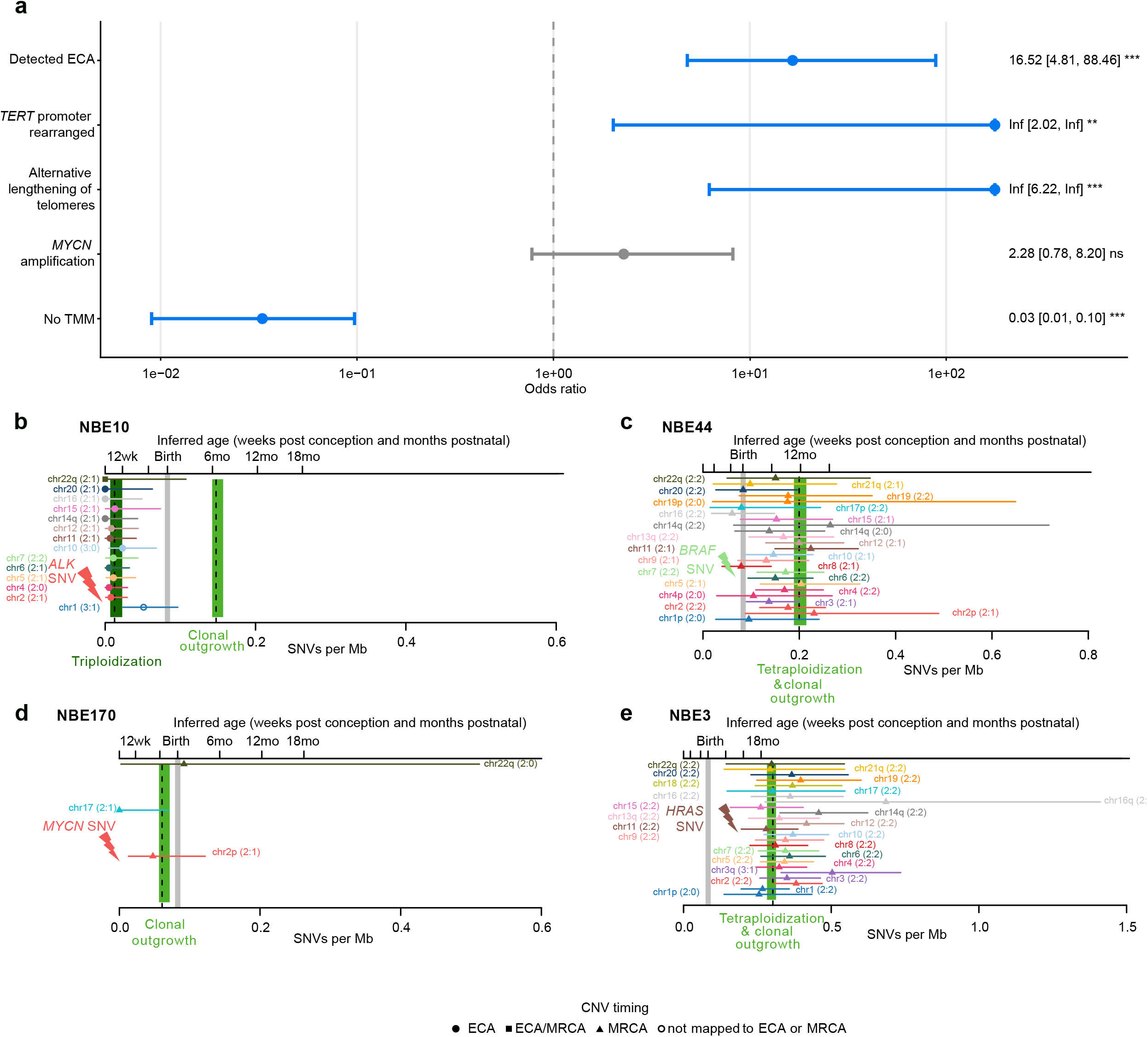
Evolution of high-risk neuroblastomas. **a**, Forest plot showing odds ratios and 95% confidence interval for associations between late MRCA tumors and detection of an ECA, as well as acquired telomere maintenance mechanisms (TERT promoter rearrangements, alternative lengthening of telomeres, *MYCN* amplification). Odds ratios were estimated using Fisher’s exact test, P values were adjusted using Benjamini-Hochberg, significant associations are highlighted in blue. **b-e**, Inferred evolutionary timelines of neuroblastomas with clonal driver mutations preceding the corresponding chromosomal gain. **b** *ALK* mutation preceding gain of Chromosome 2. **c**, *BRAF* mutation preceding gain of Chromosome 7. **d**, *MYCN* mutation preceding gain of Chromosome 2. **e**, *HRAS* mutation preceding gain of Chromosome 11. Dashed lines and shaded areas, mean and 95% confidence intervals (CIs) of estimated mutation densities at ECA and MRCA; grey area, estimated mutation density at birth. Points and error bars, mean and 95% CIs for mutation densities at chromosomal gains. Inferred age is based on the cohort-derived mutation rate of 2.17 SNVs per day (95% CI: 1.52-3.77 SNVs per day; Extended Data Fig. 1a, b).

**Extended Data Figure 3.**
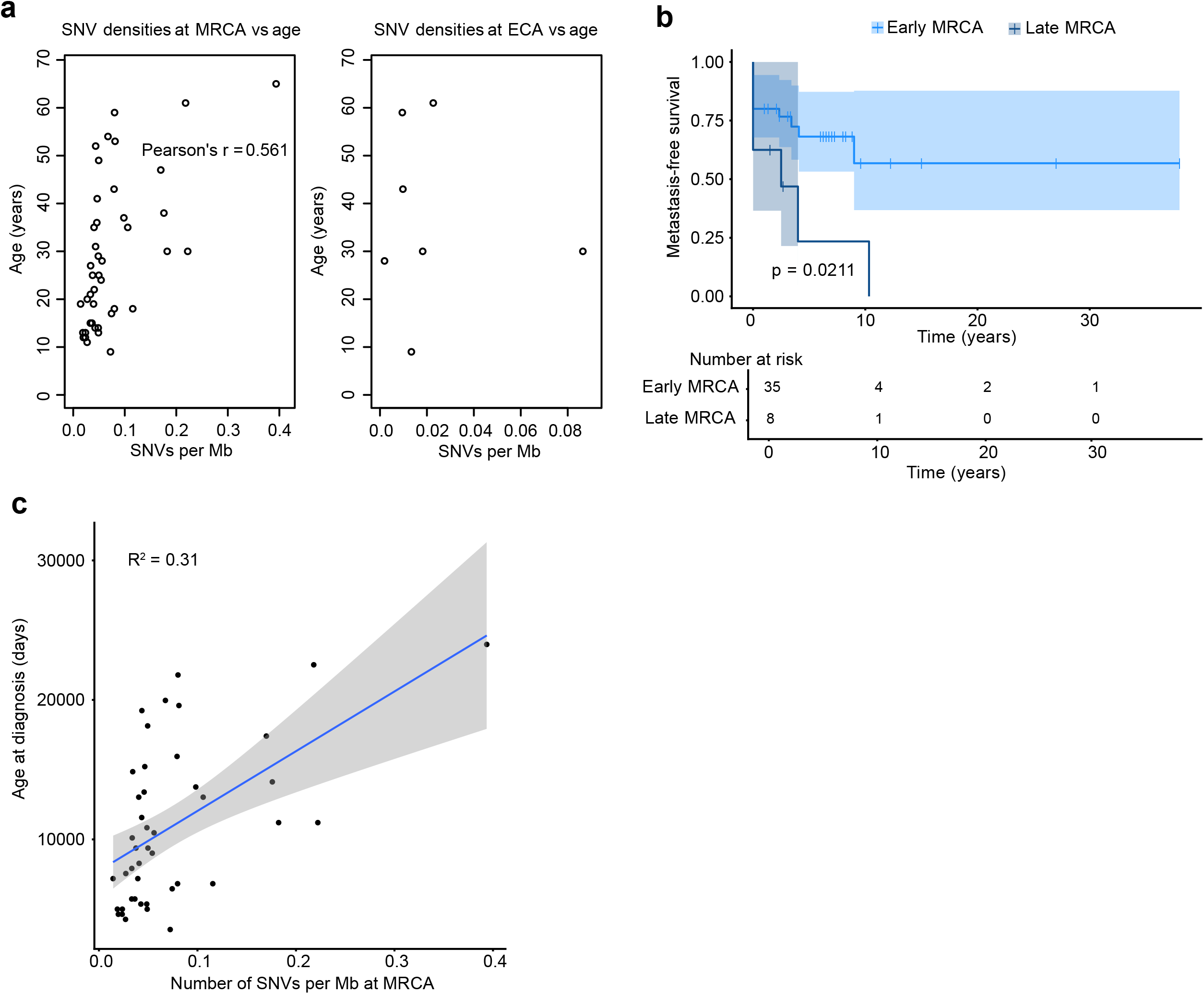
Clinical correlation of MRCA timing in paragangliomas. **a**, Correlation between age at diagnosis and mutation density at MRCA and ECA for 43 paragangliomas. **b**, Metastasis-free survival of 43 paragangliomas stratified by MRCA time. Shown are mean and 95% confidence bounds according to a log-rank test (P=0.0211). Tumors with mutation densities at MRCA below 0.1 SNVs per Mb were classified as early-MRCA. **c**, Linear regression relating age at diagnosis to the SNV burden in the tumor’s MRCA across 43 paragangliomas. Each point represents one tumor. The solid line shows the fitted linear regression (R^2^=0.31), and the grey shaded area indicates the 95% confidence interval of the fitted mean.

## Acknowledgments

We thank the members of the Westermann group and Sarah Benedetto for discussions.

## Contributions

A.M., T.H., F.W. and V.K. conceived the study. T.H. and V.K. developed the theory. M.E., A.M. and V.K. developed the code. M.E. developed the ShinyApp. M.E. and V.K. analyzed the data. M.A., V.K. and F.W. jointly supervised the study. C.B. provided additional datasets and tested the tool. M.E. and V.K. wrote the manuscript with input from all co-authors. T.H.,

F.W. and V.K. acquired funding and administered the project.

